# Comparative analysis of principal components can be misleading

**DOI:** 10.1101/007369

**Authors:** Josef C. Uyeda, Daniel S. Caetano, Matthew W. Pennell

## Abstract

Quantitative geneticists long ago recognized the value of studying evolution in a multivariate framework (Pearson, 1903). Due tolinkage, pleiotropy, coordinated selection and mutational covariance, the evolutionary response in any phenotypic trait can only be properly understood in the context of other traits (Lande, 1979; Lynch and Walsh, 1998). This is of course also well-appreciated by comparative biologists. However, unlike in quantitative genetics, most of the statistical and conceptual tools for analyzing phylogenetic comparative data (recently reviewed in Pennell and Harmon, 2013) are designed for analyzing a single trait (but see, for example Revell and Harmon, 2008; Revell and Harrison, 2008; Hohenlohe and Arnold, 2008; Revell and Collar, 2009; Schmitz and Motani, 2011; Adams, 2014b). Indeed, even classical approaches for testing for correlated evolution between two traits (e.g., Felsenstein, 1985; Grafen, 1989; Harvey and Pagel, 1991) are not actually multivariate as each trait is assumed to have evolved under a process that is independent of the state of the other (Hansen and Orzack, 2005; Hansen and Bartoszek, 2012). As a result of these limitations, researchers with multivariate datasets are often faced with a choice: analyze each trait as if they were independent or else decompose the dataset into statistically independent set of traits, such that each set can be analyzed with the univariate methods.

---

Principal components analysis (PCA) is the most common method for reducing the dimensionality of the dataset prior to analyzing the data using phylogenetic comparative methods. PCA is a projection of multivariate data onto a new coordinate system. The first PC axis is the eigenvector in the direction of greatest variance, the second PC axis, the second greatest variance, and so on. While PCA is simply another way of representing a dataset, whether or not one can draw meaningful inferences from the PC axes will depend on both the question and the structure of the data. Evolution introduces a particular kind of structure into comparative data: as a result of shared common ancestry, close relatives are likely to share many traits and trait combinations. Performing comparative analyses without considering the species’ evolutionary relationships is anathema to most evolutionary biologists, but the influence of phylogeny on data transformations is less understood (Revell, 2009; Polly et al., 2013).

Standard PCA continues to be regularly used in comparative biology. Researchers fitmod-els to PC scores computed from a variety of trait types including geometric morphometric landmarks (e.g., Dornburg et al., 2011; Hunt, 2013), measurements of multiple morphological traits (e.g., Harmon et al., 2010; Weir and Mursleen], 2013; Pienaar et al., 2013; Price et al., 2014), and climatic variables (e.g., Kozak and Wiens, 2010; Schnitzler et al., 2012). The papers we have cited here are simply examples selected from a substantial number where standard PCA was used.

The most common approach for incorporating the non-independence of species is to assume a phylogenetic model for the evolution of measured traits and use the expected covariance in the calculation of the PC axes and scores (phylogenetic principal components analysis, or pPCA; Revell, 2009). Revell’s method, explained in detail below, assumes that the measured traits have evolved under a multivariate Brownian motion (BM) process of trait evolution. Revell (2009) demonstrated that standard PCA produces eigenvalues and eigenvectors that are not phylogenetically independent.

In this paper, we first extend the argument of Revell (2009) and demonstrate how performing phylogenetic comparative analyses on standard PC axes can be positively misleading. This point has been made in other fields that deal with autocorrelated data, such as population genetics (Novembre and Stephens, 2008), ecology (Podani and Miklos, 2002), climatology (Richman, 1986) and paleobiology (Bookstein, 2012). However, the connection between these previous results and phylogenetic comparative data has not been widely appreciated and standard PCs continue to be regularly used in the field. We hope our paper helps change this practice.

Second, as stated above, Revell (2009) assumed that the measured traits had evolved under a multivariate BM process. As the pPC scores are not phylogenetically independent (Revell, 2009; also see discussion in Polly et al., 2013), one must use comparative methods to analyze them which will in turn require selecting an evolutionary model for the scores. The choice of model for the traits and the pPC scores are separate steps in the analysis (Revell, 2009). Researchers must assume a model for the evolution of the traits in order to obtain the pPC scores and then perform model-based inference on these scores. This introduces some circularity into the analysis: it seems likely that the choice of a model for the evolution of the traits will influence the apparent macroevolutionary dynamics of the resulting pPC scores. To our knowledge this effect has not been previously explored. Here we analyze simulated data to investigate whether assuming a BM model for the traits introduces systematic biases in the pPC scores when the generating model is different. We then analyze two empirical comparative datasets to understand the implications of these results for the types of data that researchers actually have; the traits in these datasets have certainly not evolved by a strict BM process.

Last, we consider the interpretation of evolutionary models fit to pPC axes and discuss the advantages and disadvantages of using pPCA compared to alternative approaches for studying multivariate evolution in a phylogenetic comparative framework. We argue that the statistical benefits of using pPC axes come at a substantial conceptual cost and that alternative techniques are likely to be much more informative for addressing many evolutionary questions.

## METHODS

### Overview of pPCA

Before describing our analyses, we briefly review standard and phylogenetic PCA and highlight the differences between the two (see Polly et al., 2013, for a more detailed treatment). In conventional PCA, a m × m covariance matrix **R** is computed from a matrix of trait values **X** for the *n* species and *m* traits 
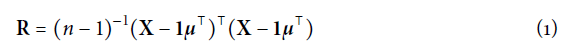
where *μ* is a vector containing the means of all *m* traits and **1** is a column vector of ones. We note that in many applications **X** may not represent the raw trait values. In geometric morphometrics for example, size, translation and rotation will often be removed from **X** prior to computing **R** (Rohlf and Slice, 1990; Bookstein, 1997). The scores **S**, the trait values of the species along the PC axes, are computed as 
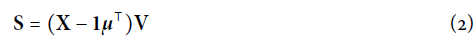
where the columns of **V** are the eigenvectors of **R**.

Phylogenetic PCA differs from this procedure in two important ways (Revell, 2009; Polly et al., 2013). First the covariance matrix is weighted by the inverse of the expected covariance of trait values between taxa under a given model **∑**. Under a BM model of trait evolution, **∑** is simply proportional to the matrix representation of the phylogenetic tree **C**, such that **∑***_i,j_*, is the shared path length between lineages *i* and *j* (Rohlf, 2001). Since only relative branch lengths matter under a multivariate BM process, we can simply set **∑** = C without loss of generality, though we note that the absolute magnitude of the eigenvalues will depend on the scale of the branch lengths. Second, the space is centered on the “phylogenetic means” **a** of the traits rather than their arithmetic means, which can be computed following Revell and Harmon (2008): 
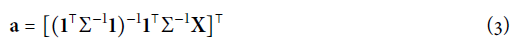

In pPCA, Equation 1 is therefore modified as 
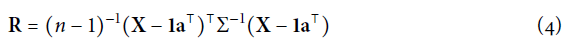

Similarly, **S** can be calculated for pPCA using Equation 2 but substituting the phylogenetic means for the arithmetic means 
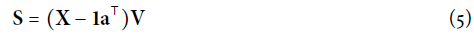
where again, **V** is a matrix containing the eigenvectors of **R**, in this case obtained from Equation 4.

The effect of weighting the covariance and centering the space using phylogeny has an important statistical consequence (Revell, 2009; Polly et al., 2013). In PCA, each PC score is independent of all other scores from the same PC axis and from scores on other axes. Due to the phylogenetic structure of the data, this property of independence does not hold when using pPCA. Therefore it is necessary to analyze pPC scores using phylogenetic comparative methods, just as one would for any other trait (Revell, 2009).

### Effect of PCA on Model Selection Under Multivariate Brownian Motion

We simulated 100 replicate datasets under multivariate Brownian motion to evaluate the effect of using standard versus phylogenetic PCA to infer the mode of evolution. For each dataset, we used TreeSim (Stadler, 2011) to simulate a phylogeny of 50 terminal taxa under a pure-birth process and scaled each tree to unit height. We then simulated a 20-trait dataset under multivariate Brownian motion. For each simulation, we generated a positive definite covariance matrix for **R**, by drawing eigenvalues from an exponential distribution with a rate λ = 1/100 and randomly oriented orthogonal eigenvectors to reflect the heterogeneity and correlation structure typical of evolutionary rate matrices (Mezey and Houle, 2005; Griswold et al., 2007). We then used this matrix to generate a covariance matrix for the tip states **X** ∼ *N*(**0**, **R** ⊗ **C**) where ⊗ denotes the Kronecker product. For each of the 100 simulated datasets, we computed PC scores using both standard methods and pPCA (using the phytools package; Revell, 2012). We used phylolm (Ho and Ane, 2014) to fit models of trait evolution to the original data and to all PC scores obtained by both methods. Following Harmon et al. (2010), we considered three models of trait evolution: 1) BM; 2) Ornstein-Uhlenbeck with a fixed root (OU: Hansen, 1997); and 3) Early Burst (EB: Blomberg et al., 2003; Harmon et al., 2010). We then calculated the Akaike Information Criterion weights (AICw) for each model/transformation/trait combination.

To explore the effect of trait correlation on inference, we conducted an additional set of simulations where **R** was varied from the above simulations to result in more or less correlated sets of phenotypic traits. We drew eigenvalues **m** from an exponential distribution and scaled these so that the leading eigenvalue **m**_1_ was equal to 1. We then exponentiated this vector across a sequence of exponents ranging for ≪1 to ≫1; this gave us a series of covariance matrices ranging from highly correlated (*m*_1_ = 1; *m*_2_,…, *m*_20_ ≈ 0) to nearly independent (**m** ≈ 1), respectively. We chose the series of exponents to obtain a regular sequence of inline1 from 0.05 to 1. For each set of eigenvalues, we simulated 25 datasets and estimated the slope of the relationship between the absolute size of phylogenetically independent contrasts (Felsenstein, 1985) and the height of the node at which they were calculated (the “node height test”; Purvis and Rambaut, 1995). Under OU models, this relationship is expected to be positive, while under EB models this relationship is negative. BM models are expected not to show correlation between contrasts and height of the nodes.

### Effect of Using PCA When Traits are not Brownian

We then simulated datasets under alternative models of trait evolution. First, we also simulated traits under a correlated multivariate OU model using the mvSLOUCH package (Bartoszek et al., 2012). Combined with the correlated BM simulations above, we used correlated OU simulations to explore the effect of PCA and pPCA on model inference under reasonably biologically realistic conditions. We simulated 20 correlated traits for 50 taxa trees using a positive definite covariance matrix for the diffusion matrix by drawing eigenvalues from an exponential distribution with a rate λ = 1 and randomly oriented orthogonal eigenvectors. The a-matrix was set a diagonal matrix with a constant value of 2 for each trait such that the phylogenetic half-life log(2)/α (Hansen et al., 2008) was approximately equal to 0.35 of the total tree depth. The root state for each simulation was set at the multivariate phenotypic optimum. We then fit BM, OU and EB models to the original data, PC scores and pPC scores for each simulated dataset and estimated parameters and AICw.

Second, we simulated an additional set of datasets with uncorrelated traits and equal evolutionary rates. These simplified datasets allowed us to generate comparable data under all three generating models (BM, OU and EB) and isolate how misspecifying the model of trait evolution can impact the distribution of PC and pPC scores. As before, for each model we simulated 20 traits on 50 taxa trees. For the BM simulations, we set *σ*^2^ = 1 for all 20 traits. For OU, we set *σ*^2^ = 1 and *α* = 2. For EB, we again set *σ*^2^ = 1 and set *r*, the exponential rate of deceleration, to be log(0.02). As above, we estimated parameters and AICw for each model fit to original data, PC scores and pPC scores. In addition, we applied two common diagnostic tests for deviation from BM-like evolution to all trait/PC axes. First, we calculated the slope of the node height test as described in the preceding section. Second, we characterized the disparity through time (Harmon et al., 2003) using the geiger package (Pennell et al., 2014a).

Finally, we examined the scenario in which a set of traits each follow a model of evolution with unique evolutionary parameters. In particular, we use the accelerating-decelerating (ACDC) model of Blomberg et al. (2003) to generate independent trait datasets. This model is a general case of the EB model which allows both accelerating or decelerating rates of phenotypic evolution. Accelerating rates of evolution result in identical likelihoods as the OU model (assuming the root state is at the optimal trait value and the tree is ultrametric), and thus are equivalent for our purposes (we provide a proof for this claim in the Supplementary Material). We simulated 100 datasets with 50 taxa and 20 traits. Trees were generated as in previous simulations. Each trait was simulated along the phylogeny with an exponential rate of change *r* drawn from a normal distribution with mean 0 and standard deviation of 5. Values of *r* above 0 correspond to accelerating evolutionary rates, while those below 0 correspond to decelerating, or Early-Burst models of evolution. For each dataset, we conducted both standard and phylogenetic PCA in which the traits are standardized to unit variance (i.e., using correlation matrices, which ensured traits across parameter values had equal expected variances). For each PC or pPC, we regressed the magnitude of the trait loadings against the trait’s ACDC parameter value. We then visualized whether there were systematic trends in the relationship between the ACDC parameter value, and the weight given to a particular trait across PC axes. Such systematic trends would indicate that either PCA or pPCA “sorts” traits into PC axes according to the particular evolutionary model each trait follows.

### Empirical Examples

We analyzed two comparative datasets assembled from the literature, allowing us to investigate the effects of principal components analyses on realistically structured data. First, we analyzed phenotypic evolution across the family Felidae (cats) using measurements from two independent sources—five cranial measurements from Slater and Van Valkenburgh (2009) and body mass and skull width from Sakamoto et al. (2010). For the analysis, we used the supertree compiled by Nyakatura and Bininda-Emonds (2012). Second, we analyzed 23 morphometric traits in *Anolis* lizards and phylogeny from Mahler et al. (2010). In both datasets, all measurements were linear measurements on the logarithmic scale. We conducted standard and phylogenetic PCA and examined the effect of each on model-fitting, the slope of the node height test, and the average disparity through time. All simulations and analyses were conducting using R v3.0.2. Scripts to reproduce our results are available at https://github.com/mwpennell/phyloPCA.

## RESULTS

### Effect of PCA on Model Selection Under Multivariate Brownian Motion

Standard PCA introduces a systematic bias in the favored model across principal components. In our simulations where the traits evolved under a multivariate BM model, EB models had systematically elevated support as measured by Akaike weights for the first few components, for which it generally exceeded support for the BM model (Figure 1, left panel). Fitting models sequentially across PC axes 1−20 revealed a regular pattern of increasing support for BM models moving from the first toward the intermediate components, followed by increasing support for OU models among later components, which generally approached an AICw of 1. This regular pattern across trait axes was not present for either the original datasets, or for phylogenetic principal components, which found strong support for the BM model regardless of which trait was analyzed. As BM is a special case of both OU and EB, the likelihoods for the more complex models will converge on that of BM when the true model is Brownian. AIC weights for model *i* are computed as AICw_i_ = exp[0.5(AIC*_min_* − AIC*_i_*)]/ Σ*_j_* exp[0.5(AIC*_min_* − AIC*_j_*)] and therefore if the likelihoods are identical, OU and EB will have a ΔAIC = AIC*_min_* − AIC*_i_* = 2 (as OU and EB each have one more parameter than BM). The theoretical maximum for the AICw of BM is thus 1/(2e^−1^+ 1) ≈ 0.576.

**Figure 1:**
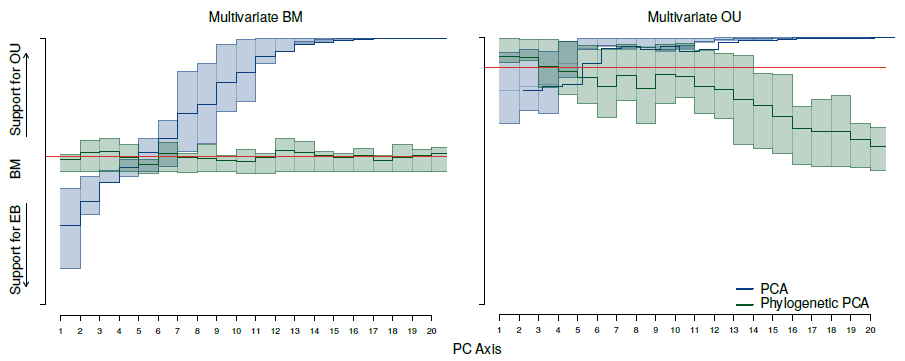
Distribution of support for BM, OU and EB models when the generating model is a correlated multivariate BM model (left panel) and OU model (right panel). Support for models were transformed onto a linear scale by calculating an overall model support statistic: *AlCw*_OU_* – AICw*_EB_. Thus high values support OU, low values support EB, and intermediate values near 0 indicate BM-like evolution. Models were fit to each replicated dataset for each of 20 different traits which were taken either from PC scores (blue line) or phylogenetic PC scores (green line). Shaded regions indicate the 25^th^ and 75^th^ quantiles of the model support statistic for 100 replicated datasets. The red line indicates the average model support statistic averaged over all 20 original trait variables.

Multivariate datasets simulated with high correlations (i.e., low effective dimensionality) showed increased support for BM across PC axes. When the leading eigenvalue explained a large proportion of the variance, the slope of the node height test converged toward 0, indicating no systematic distortion of the contrasts through time (Figure 2). However, when the eigenvalues of the rate matrix were more even, standard PCA resulted in a negative slope in the node height test among the first few PCs, which in turn provides elevated support for EB models. This pattern is reversed among higher PC axes, which have positive slopes between node height and absolute contrast size, which provides elevated support for OU models (Figures 2 and 3).

**Figure 2:**
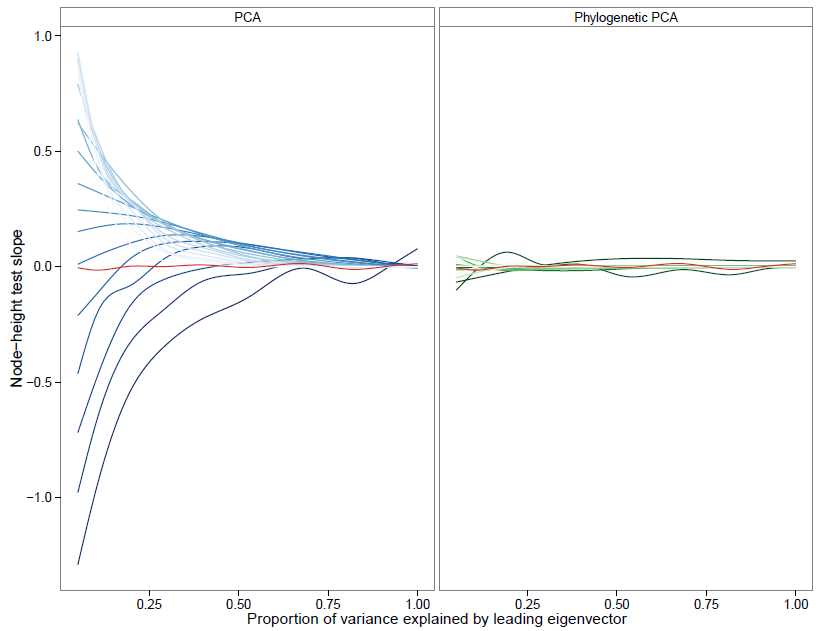
Effect of trait correlations on the slope of the node height test for PC scores (left) and pPC scores (right) under a multivariate BM model of evolution. The red line is the aggregated data for all 20 traits on the original (untransformed) scale. The intensity of the colors are proportional to the ranking of the PC or pPC axes, stronger lines represent the first axes. When the leading eigenvector explains very little variation in the data and the effective dimensionality is high, the slope of node height test increases from negative to positive across PC axes. This indicates that under standard PCA, PC1 has higher contrasts near the root of the tree, while later PCs have higher contrasts near the tips (resulting in the pattern of model support observed in Figure 1). As the amount of variance explained by the principal eigenvector increases, the slope of the node height test approaches 0. No such effect is found for phylogenetic PCA.

**Figure 3:**
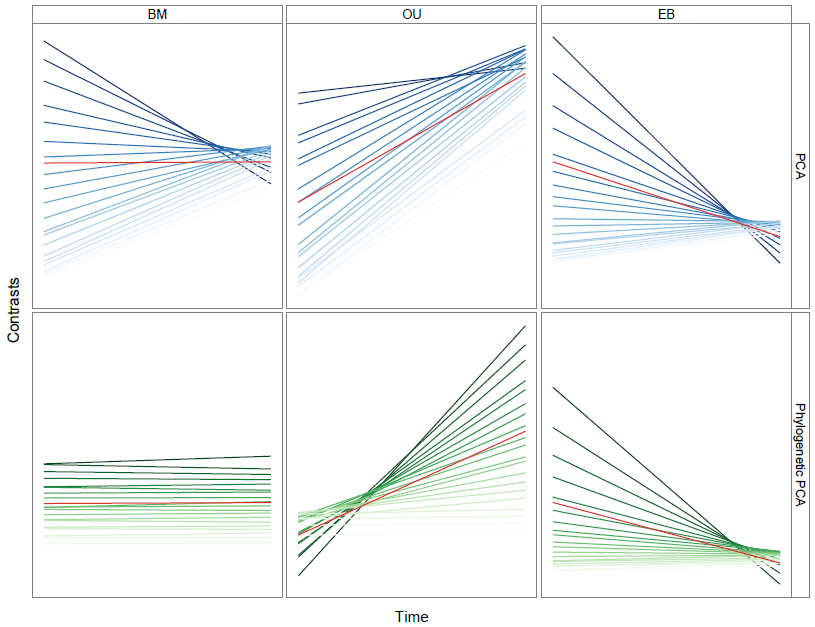
Relationship between the average phylogenetic independent contrasts and the height of the node across 100 datasets simulated under either a BM (left), OU (middle) or EB (right) model of evolution. Contrasts were calculated for each of the 20 traits corresponding to either PC scores (top row) or pPC scores (bottom row). Each line represents a best-fit linear model to the aggregated data across all 100 replicate simulations. Red lines are aggregated over all 20 traits on the original data. The plots are oriented so that the left side of each panel corresponds to the root of the phylogeny, with time increasing tipward to the right. The intensity of the colors are proportional to the ranking of the PC or pPC axes, stronger lines represent the first axes. PCA results in a predictable pattern of increasing slope in the contrasts across PCs. By contrast, pPCA only has systematic distortions across pPC axes when the underlying model is not multivariate BM. When this occurs, the first few pPC axes tend to have more extreme slopes than the original data (but in the correct direction).

### Effect of using PCA When Traits are not Brownian

If the underlying model was either OU or EB rather than BM, then phylogenetic PCA tended to increase support for the true model relative to the original trait variables for the first few component axes (Figures 1, right panel; S.1; and S.2). For example, when each of the original trait variables were simulated under a correlated or uncorrelated OU process, support for the OU model increased for pPC1 relative to the original trait variables. Higher principal component axes showed a regular pattern of decreasing support for the OU model, while the last few PCs have equivocal support for either a BM or OU model (Figures 1, right panel and S.1). Furthermore, parameter estimation was affected by phylogenetic PCA. The *α* parameter of the correlated and uncorrelated OU models were estimated to be stronger than the value simulated for individual traits for the first few pPC scores and lower for the higher components (Figure S.3 and S.4).

Examining the outcomes of the node height tests (Figure 3) and the disparity through time analyses (Figure S.5) for uncorrelated OU, EB and BM models helps clarify the results we observed from model comparison and parameter estimation. Under OU models, traits are expected to have the highest contrasts near the tips, whereas under EB models, traits will have the highest contrasts near the root of the tree. Under multivariate BM, standard PCA maximizes the overall variance explained across the entire dataset, thereby tending to select linear combinations of traits that maximize the contrasts at the root of the tree. Thus, the first few PCs are skewed toward resembling EB models, while the last few PCs are skewed toward resembling OU models. By contrast, the effect of pPCA on the node height relationship depends on the generating model. When traits are evolved under an OU model, the first few pPC axes show an exaggerated pattern of high variance towards the tips. Likewise, when traits are evolved under an EB model, the first few pPC axes show an exaggerated pattern of high variance towards the root of the tree. For traits generated under both OU and EB models, the higher components resemble BM-like patterns.

When the data includes traits evolved under ACDC models with varying parameters, both PCA and pPCA systematically assigned traits to particular PCA axes according to the parameter values of the generating model. Traits which follow EB models are preferentially given higher loadings for the first few PCs as well as the last few PCs (Figure 4). Intermediate PCs had relatively even loadings slightly skewed toward accelerating rates (i.e., OU-like models), while most of the traits with decelerating rates were assigned with high loadings to just a few PC axes. This asymmetry may reflect the fact that EB models are more variable in their outcomes to the phylogeny, owing to the fewer independent branches among which divergence can occur closer to the root of the tree. Our results indicate that both pPCA and PCA can be biased in the selection of PC axes with respect to the generating evolutionary model.

**Figure 4:**
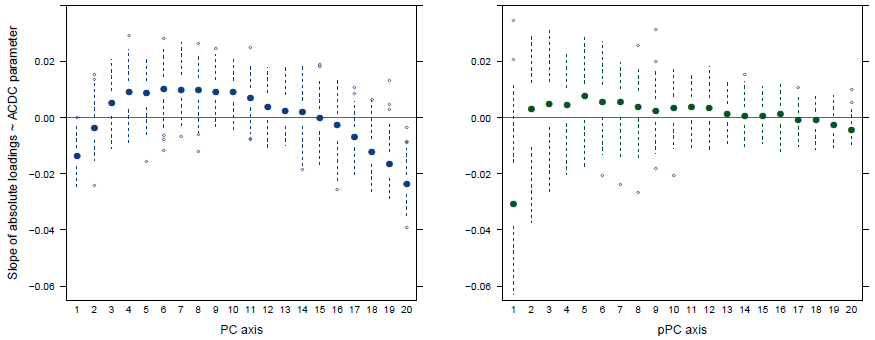
Relationship between factor loadings and ACDC parameter (α) for PCA (left) and pPCA (right) across 100 simulated datasets. For each simulation a value of α were drawn from a Normal distribution with mean = 0 and sd = 5. Boxplots indicate the distribution of the slope of a linear model describing the relationship between the absolute factor loadings for a given PC and the magnitude of the ACDC parameter. A negative slope indicates that traits with decelerating rates of evolution tend to have higher loadings in that particular PC. Conversely, positive slopes indicate that traits with accelerating rates tend to have higher loadings.

### Empirical Examples

In the felid dataset, the seven morphometric traits were extremely highly correlated, with the first PC explaining 96.9% and 93.7% of the total variation in the dataset for standard PCA and phylogenetic PCA, respectively. All raw traits and the first PC axis of both standard and phylogenetic PCA support a BM model of evolution (PC and pPC axes have AICw’s of 0.574, which is near the theoretical maximum for BM). The last four standard PC axes show strong support for an OU model (AICw ≈ 1) whereas under phylogenetic PCA the last axes have mixed support favouring BM or OU (Figure S.6). Both the node height test and the disparity through time plots show this same pattern. The node height slope of the first axis is approximately zero while the slope of the remaining axes are slightly positive under standard and phylogenetic PCA. The first axis show the same disparity through time pattern of the untransformed data in both standard and phylogenetic PCA. However, the last PC axes show disparity accumulated toward the tips under standard PC, while phylogenetic PCA produced a less clear pattern (Figure S.7).

For the morphometric traits in the *Anolis* dataset, the first PC also explained a large proportion of the variation (92.6% and 90.0% for standard and phylogenetic PCA, respectively). Most of the untransformed traits had equivocal support for either a BM or EB model (Figure S.8). While PC1 of both PCA and pPCA mirrored this pattern, the remaining PCs for both PCA and pPCA show a general pattern of decreasing support for an EB model (Figure S.8). Collectively PCs2−4 had higher support for the EB model than any other PC in both standard PCA (AICw*_EB_*: PC2 = 1.0; PC3 = 0.47; PC4 = 0.28) and phylogenetic PCA (AICw*_EB_*: pPC2 = 1.0; pPC3 = 0.43, pPC4 = 0.27). Similarly, these early PC axes tended to have more negative slopes from the node height test relative to the average trait in the dataset (Figure S.9).

## DISCUSSION

Different ways of representing the same set of data can change the meaning of measurements and alter the interpretations of subsequent statistical analyses (Houle et al., 2011). PCA is often considered to be a simple linear transformation of a multivariate dataset and the potential consequences of performing phylogenetic comparative analyses on PC scores have received very little attention. In this paper, we sought to highlight the fact that fitting macroevolutionary models to a handful of PC axes may positively mislead inference—what appears like the signal of an interesting biological process may simply be an artifact stemming from how PCA is computed. By focusing analyses exclusively on the first few PC axes, as is commonly done in comparative studies, researchers are, in effect, taking a biased sample of a multivariate distribution (Mitteroecker et al., 2004). We demonstrate how this biased sampling can affect inferences from both PCA and pPCA. In particular, we demonstrate that it can lead researchers to erroneously infer a pattern of decreasing rates of evolution through time in highly dimensional datasets.

We can obtain an intuitive understanding of how PCA can affect inferences by considering data simulated under a multivariate BM model. Despite a constant rate of evolution across each dimension of trait space, stochasticity will ensure that some dimensions will diverge more rapidly than expected early in the phylogeny, while others will diverge less. All else being equal, dimensions that happen to diverge early are expected to have the greatest variance across species and standard PCA will identify these axes as the primary axes of variation. However, the trait combinations that are most divergent early in the clade will appear to have slowed down towards the present simply due to regression toward the mean, resulting in the characteristic “early burst” pattern of evolution for the first few principal components. (Pennell et al. 2012 pointed out that lineage diversification models are susceptible to a similar sampling effect.) An analogous process will result in the last few PCs following an OU process, in which the amount of divergence will be concentrated toward the present. Standard PCA thus effectively “sorts” orthogonal trait dimensions by whether they follow EB, BM and finally, OU like patterns of trait divergence. Traits studied using PCA may therefore often be biased to reflect particular evolutionary models, merely as a statistical artifact.

These problems ultimately stem from making statistical inferences from a selected few PC axes without accounting for how PCA transforms autocorrelated data. This issue is certainly not limited to phylogenetic comparative studies (see Richman, 1986; Podani and Miklos, 2002; Jolliffe, 2002; Novembre and Stephens, 2008; Bookstein, 2012). For example, Novembre and Stephens (2008) demonstrated that apparent waves of human migration in Europe obtained from PCA of genetic data (e.g.,Cavalli-Sforza et al., 1994) could be attributed to artifacts similar to those we document here; in their case, the auto-correlation was the result of geography rather than phylogeny. While the bias introduced by analyzing standard PCs with phylogenetic models has been documented previously (Revell, 2009; Polly et al., 2013), we sought to clarify precisely how inferences of macroevolutionary processes and patterns can be impacted.

Revell (2009) recognized the need for accounting for phylogenetic correlation when performing PCA transformations and introduced the phylogenetic PCA method. Our simulations verify that when the underlying model is multivariate BM, pPCA mitigates the effect of deep divergences among clades in the major axes of variation by scaling divergence by the expected divergence given the branch lengths of the phylogeny. However, BM is often a poor descriptor of the macroevolutionary dynamics of trait evolution (for example, see Harmon et al., 2010; Pennell et al., 2014b) and assuming this model when performing pPCA is less than ideal. Revell (2009) suggested that alternative covariance structures could be used to estimate phylogenetically independent PCs for different models. For example, one could first optimize the λ model (Freckleton et al., 2002) across all traits simultaneously and then rescale the branch lengths of the tree according to the estimated parameter in order to obtain Σ for use in Equation 4. However, one cannot compare model fits across alternative linear combinations of traits, so the data transformation must occur separately from modeling the evolution of the PC axes. As Revell (2009) noted, parameters estimated to construct the co variance structure for the pPCA will likely be different from the same parameters estimated using the PC scores themselves. Furthermore, this procedure is restricted to models that assume a shared mean and variance structure across traits (see Hansen et al., 2008; Hansen and Bartoszek, 2012; Bartoszek et al., 2012, for examples where this does not apply). As such, if the question of interest relies on model-based inference, transforming the data using pPCA necessitates *ad hoc* assumptions about the evolution of the traits, and researchers must hope that the resulting inferences are generally robust to these decisions.

We show that when the trait model is misspecified, systematic and predictable distortions occur across pPC axes—similar to those that were observed when the phylogeny was ignored altogether. In some scenarios such distortions may not substantially alter inference. For example, when all traits evolve under an OU model (or when all traits evolve under a EB model), we find that these distortions primarily serve to inflate the support for the true model. Even so, interpretation of parameter estimates for pPC scores becomes much more challenging (Figures 3, S.3, S.4, and S.5). More complex scenarios can produce more worrying distortions. When evolutionary rates vary through time and across traits, both PCA and pPCA will sort traits into PC axes according to which evolutionary model they follow, despite all traits being evolutionarily independent. Under the conditions we examined, this resulted in both PC1 and pPC1 being heavily-weighted toward EB-type models, despite simulating an even distribution of accelerating and decelerating rates across traits. Intriguingly, we observe similar patterns for both PCA and pPCA in the *Anolis* morphometric dataset (Figures S.8 and S.9). Focusing on the first few axes of variation identified by pPCA alone may skew our view of evolutionary processes in nature, and bias researchers toward finding particular patterns of evolution.

When employed as a descriptive tool, PCA can be broadly used even when assumptions regarding statistical non-independence or multivariate normality are violated (Jolliffe, 2002). There is nothing wrong with using standard PCA or pPCA on comparative data to describe axes of maximal variation across species or for visualizing divergence across phylo-morphospace (Sidlauskas, 2008). Furthermore, our simulations and empirical analyses suggest that strong correlations among traits (i.e., when the leading eigenvector explained a majority of the variation, e.g., > 75%), PC scores may not be appreciably distorted (Figure 2). The statistical artifacts we describe are more likely to appear when matrices have high effective dimensionality (see Bookstein, 2012). Given that many morphometric datasets may be highly correlated, the overall effect of using standard PCA or of misspecifying the model in phylogenetic PCA may in some cases be relatively benign.

And we certainly do not mean to imply that the biological inferences that have been made from analyzing standard or phylogenetic PC scores in a comparative framework are necessarily incorrect. When Harmon et al. (2010) analyzed the evolution of PC2 (what they referred to as “shape”) obtained using standard PCA, they found very little support for the EB model across their 39 datasets. The magnitude of the bias introduced by using standard PCA is difficult to assess but any bias that did exist would be towards finding EB-like patterns. This only serves to strengthen their overall conclusion that such slowdowns are indeed rare (but see Slater and Pennell, 2014). Our results do suggest that in some cases analyses conducted with PC axes should be revisited to ensure that results are robust.

The broader question raised by our study is how one should draw evolutionary inferences from multivariate data. The first principal component axis from pPCA is the phylogenetically-weighted “line of divergence”, the major axis of divergence across the sampled lineages in the clade (Hohenlohe and Arnold, 2008). This axis is of considerable interest in evolutionary biology. The direction of this line of divergence may be affected by the orientation of within-population additive genetic (co)variance **G**, such that evolutionary trajectories may be biased along the “genetic line of least resistance”; i.e., divergence occurs primarily along the leading eigenvector of **G**, **g**_max_ (Schluter, 1996). Alternatively, the line of divergence may align with ω_max_, the “selective line of least resistance”, due to the structure of phenotypic adaptive landscapes (Arnold et al., 2001; Jones et al., 2007; Arnold et al., 2008), or else may be driven by patterns of gene flow between populations (Guillaume and Whitlock, 2007) or the pleiotropic effects of new mutations (Jones et al., 2007; Hether and Hohenlohe, 2014).

While it is perfectly sensible and interesting to compare the orientation of pPC1 to that of **g**_max_, ω_max_ or other within-population parameters, making explicit connections between macro-and microevolution requires a truly multivariate approach. Quantitative genetic theory makes predictions about the overall pattern of evolution in multivariate space (Lande, 1979). By fitting evolutionary models to pPC scores, we are only considering evolution along these axes independently and not fully addressing potentially relevant patterns in the data. In contrast, multivariate tests for the correspondence of axes of trait variation within and between species can provide meaningful insights into the processes by which traits evolve over long time scales (Hohenlohe and Arnold, 2008; Bolstad et al., 2014).

The most conceptually straightforward multivariate approach for analyzing comparative data is to construct models in which there is a covariance in trait values between species (which is done in univariate models) and a covariance between different traits. Such multivariate extensions of common quantitative trait models have been developed (Butler and King, 2004; Revell and Harmon, 2008; Hohenlohe and Arnold, 2008; Revell and Collar, 2009; Thomas and Freckleton, 2012). These allow researchers to investigate the connections between lines of divergence and within-population evolutionary parameters (Hohenlohe and Arnold, 2008) as well as to study how the correlation structure between traits itself changes across the phylogeny (Revell and Collar, 2009).

These approaches also have substantial drawbacks. First, the number of free parameters of the models rapidly increases as more traits are added (Revell and Harmon, 2008), making them impractical for large multivariate datasets. This issue may be addressed by constraining the model in meaningful ways (Butler and King, 2004; Revell and Collar, 2009) or by assuming that all traits (or a set of traits) share the same covariance structure (Klingenberg and Marugan-Lobon, 2013; Adams, 2014b,a). Such restrictions of parameter space are especially appropriate for truly high-dimensional traits, such as shape. For such traits, we are primarily interested in the evolution of the aggregate trait and not necessarily the individual components (Adams, 2014b). The second drawback is that these models allow for inference of the covariance between traits but the cause of this covariance is usually not tied to specific evolutionary processes. This difficulty can be addressed by explicitly modeling the evolution of some traits as a response to evolution of others. Hansen and colleagues have developed a number of models in which a predictor variable evolves via some process and a response variable tracks the evolution of the first as OU process (Hansen et al., 2008; Hansen and Bartoszek, 2012; Bartoszek et al., 2012). This has been a particularly useful way of modeling the evolution of allometries (e.g., Hansen and Bartoszek, 2012; Voje et al., 2013; Bolstad et al., 2014). But, as with the multivariate versions of standard models discussed above, increasing the number of traits makes the model much more complex and parameter estimation difficult.

As we can only estimate a limited number of parameters from most comparative datasets— and even when we consider large datasets, most existing comparative methods have only been developed for the univariate case—it often remains necessary to reduce the dimensionality of a multivariate dataset to one or a few compound traits. We believe that although PCA can be potentially quite usefully applied to this problem, it may be in ways that are statistically and conceptually distinct from how it is conventionally applied to comparative data.

First, we argue that reducing multivariate problems to more easily managed, lower dimensionality analyses should be approached with the specific goal of maintaining biological meaning and interpretability (Houle et al., 2011). The common practice of examining only the first few PCs carries with it the implicit assumption that PCA ranks traits by their evolutionary importance, though this is not necessarily true (Polly et al., 2013). If a certain PC axis is of sufficient biological interest in its own right, it may not matter if it is a biased subset of a multivariate distribution. The fact that a vast majority of the traits studied in adaptive radiations likely represent very biased axes of variation across the multivariate process of evolution does not diminish the importance of the inferences made from studying these traits.

The danger occurs when the biological significance of the set of traits is poorly understood, and when the source of the statistical signal maybe either artifactual or biological. If a trait was not of interest *a priori,* then this essentially turns into a multiple comparisons problem in which PCA searches multivariate trait space for an unusual axis of variation. These axes will tend to appear to have evolved by a process inconsistent with the generating multivariate process as a whole. *A posteriori* interpretation of the PC axes by their loadings is something of an art—one must “read the tea leaves” to understand what these axes mean biologically. Even when a particular axis is correlated with a biological interpretation, it can be unclear whether the statistical signal supporting a particular inference results from the evolutionary dynamics of the trait of interest or if it is the result of statistical artifacts introduced by the imperfect representation of that trait by a PC axis. More rigorous algorithms can be applied to identify subsets of the original variables that best approximate the principal components, which although still biased, are frequently more interpretable (Hausman, 1982; Somers, 1986,1989; Vines, 2000; Cadima and Jolliffe, 2001; Jolliffe, 2002; Zou et al., 2006). Another potential approach is to use principal components computed from within-population data, rather than comparative data. For example, if **G**, or failing that, the phenotypic variance-covariance matrix **P**, is available for a focal species, then the traits associated to the principal axes of variation in that species can be measured across all species in the phylogeny. In other words, across species trait measurements can be projected along **g**_max_. This alleviates the issues we discuss in this paper by estimating PCs from within-population data that is independent from the comparative data used for model-based inference.

Components defined by within-population variance structure or by approximating principal components with interpretable linear combinations will not explain as much variance across taxa as standard PCA and will not necessarily be statistically independent of one another. But the extra variance explained by the principal components of comparative data may in fact include a sizeable amount of stochastic noise, rather than interesting biological trait variation, as we have shown in our simulations. Furthermore, while the trait combinations (eigenvectors) identified by pPCA will be statistically orthogonal, this is only true in the particular snapshot captured by comparative data and does not imply that they are evolving independently. The distinction between statistical and evolutionary independence is crucial (Hansen and Houle, 2008) but it is easy to conflate these concepts when the data has been abstracted from its original form. We argue that the added intepretability of carefully chosen and biologically meaningful trait combinations far outweighs the cost of some trait correlations or explaining less-than-maximal variation.

## CONCLUDING REMARKS

In this note we sought to clarify some statistical and conceptual issues regarding the use of principal components in comparative biology. We have shown that fitting evolutionary models to standard PC axes can be postively misleading. And despite the development of methods to correct for this, in our reading of the empirical literature, we have found this to be a common oversight. We have also demonstrated that misspecifying the model of trait evolution when conducting pPCA may influence the inferences we make from the pPC scores. We show that in some scenarios, pPCA may sort traits according to the particular evolutionary models they follow in a similar manner as standard PCA. Ignoring phylogeny altogether is, of course, a form of model misspecification. Consequently, we caution that the use of pPCA may bias inference toward identifying particular evolutionary patterns, which may not be representative of the true multivariate process shaping trait diversification. We hope that our paper provokes discussion about how we should go about analyzing multivariate comparative data. We certainly do not have the answers but argue there are some major theoretical limitations inherent in using PCA, phylogenetic or not, to study macroevolutionary patterns and processes.

## Acknowledgements

We would like to thank our advisor, Luke Harmon, for encouraging us to pursue this project and for providing insightful comments on the work and manuscript. We thank Luke Mahler for providing data for the *Anolis* empirical example. We thank Frank Andersen, Tanja Stadler, David Polly, and two anonymous reviewers for helpful comments on this paper. JCU was supported by NSF DEB 1208912 and DBI 0939454. DSC was supported by a fellowship from Coordenação de Aperfeiçoamento de Pessoal de Nivel Superior (CAPES: 1093/12−6). MWP was supported by a NSERC postgraduate fellowship.

## APPENDIX EQUIVALENCY BETWEEN ORNSTEIN-UHLENBECK AND ACCELERATING CHANGE MODELS

In our paper, we investigate the scenario in which the individual traits have each evolved under a different model. To simulate the data, we drew values for the exponential rate parameter r of the accelerating/decelerating change (ACDC; Blomberg et al., 2003) model for each trait from a normal distribution with mean 0. We claim that when r is positive, the ACDC model generates traits with a structure equivalent to those produced by a single optimum Ornstein-Uhlenbeck (OU; Hansen, 1997) model. To our knowledge, this has not been previously demonstrated in the literature. Slater et al. (2012) suggested that these two models were equivalent for ultrametric trees: “Looking at extant taxa only, the outcome of [a process with accelerating rates] is very similar to an OU process, as both tend to erase phylogenetic signal” [p. 3940], though they did not provide any proof.

*Conjecture.* A single optimum OU process produces identical covariance matrices to those produced by the AC model when i) the tree is ultrametric and ii) the trait is assumed to be at the optimum at the root of the tree.

*Proof.* Consider a bifurcating tree of depth *T* with two terminal taxa *i* and *j* that are sampled at the present and share a common ancestor at time *t_ij_* where *t_ij_ < T*. A trait *Y* is measured for both *i* and *j*.

## ORNSTEIN-UHLENBECK PROCESS

First, assume that *Y* has evolved according to an Ornstein-Uhlenbeck (OU) process 
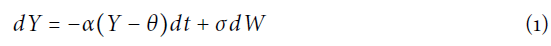
where *θ* is the optimum trait value, *α* is the strength of the pull towards *θ*, and *σ* is the rate of the Brownian diffusion process *dW* (Hansen, 1997). Also assume that the processbegan at the optimum, such that *Y(t* = 0) = *θ*. The expected value for *Y_i_* and *Y_j_* is equal to the root state. The expected variance for both *Y_i_* and *Y_j_* is given by Hansen (1997):

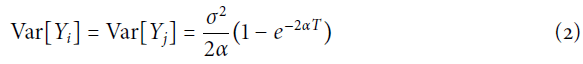

The expected covariance between lineages *Y_i_* and *Y_j_* is given by 
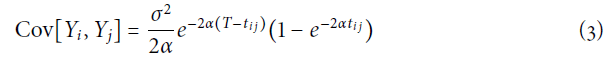

The correlation between *Y_i_* and *Y_j_*, *ρ*[*Y_i_, Y_j_*], is defined as 
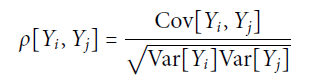

Under an OU process, *ρ*[*Y_i_, Y_j_*] is 
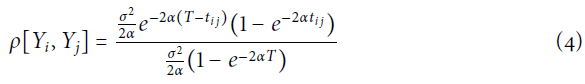

With some algebra, it is straightforward to reduce Equation 4 to 
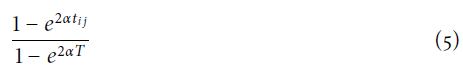

## ACCELERATING CHANGE MODEL

Next, assume that *Y* has evolved according to the Accelerating Change (AC) model, which describes a Brownian motion process in which the rate of diffusion *σ*^2^ changes as function of time 
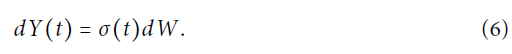

Specifically, we consider the functional form of *σ^2^(t)* to be 
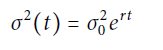

where *r* is constrained to be positive (Blomberg et al., 2003; Slater et al., 2012). The expected value of the AC model is also equal to the root state. The expected variance for *Y_i_* and *Y_j_* is given by

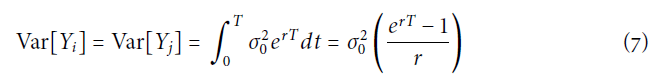

(Harmon et al., 2010) and the covariance is equal to

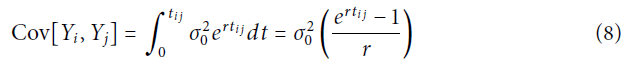

Under the AC model, *ρ*[*Y_i_, Y_j_*] is 
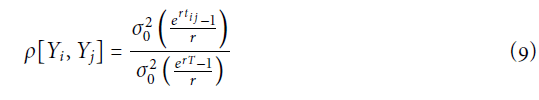

Equation 9 can be easily reduced to

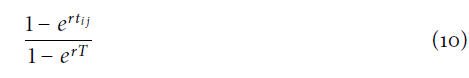

## COMPARING THE EXPECTATIONS UNDER OU AND AC

Comparing equations 4 and 9, it is clear that the correlation between *Y_i_* and *Y_j_* under the OU model is equal to that of the AC model 
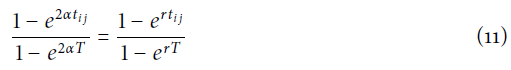
when *α* = 0.5 *r*. For every value of *α* there is a value of *r* that can produce an identical correlation structure. Note that the values of *σ*^2^ and 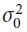 do not affect the correlation structure but do enter into the covariance structure. AsCov[*Y_i_, Y_j_*] *=* Var[*Y_i_, Y_j_*]ρ[*Y_i_, Y_j_*] and the values of *ρ*[*Y_i_, Y_j_*] are equivalent, we can set the variances of two models (given by Equations 2 and 7) equal to one another 
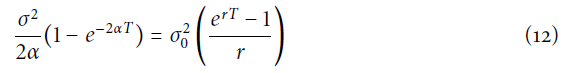
and substitute *r* for 2*α* 
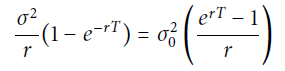

Reducing algebraically, it is easy to show that 
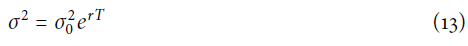

Therefore for any covariance matrix for *Y_i_ and Y_j_*, OU and AC are completely unidentifiable and the likelihoods for the two models will be identical.

*Notes.* The two variances Var[*Y_i_*] and Var[*Y_j_*] will only be equal to one another when the tree is ultrametric. If either *i* or *j* were not sampled at the present (e.g., if one was an extinct lineage), this proof for the non-identifiability of OU and AC does not hold and one can potentially distinguish these models (Slater et al., 2012).

**FIGURE S.1:**
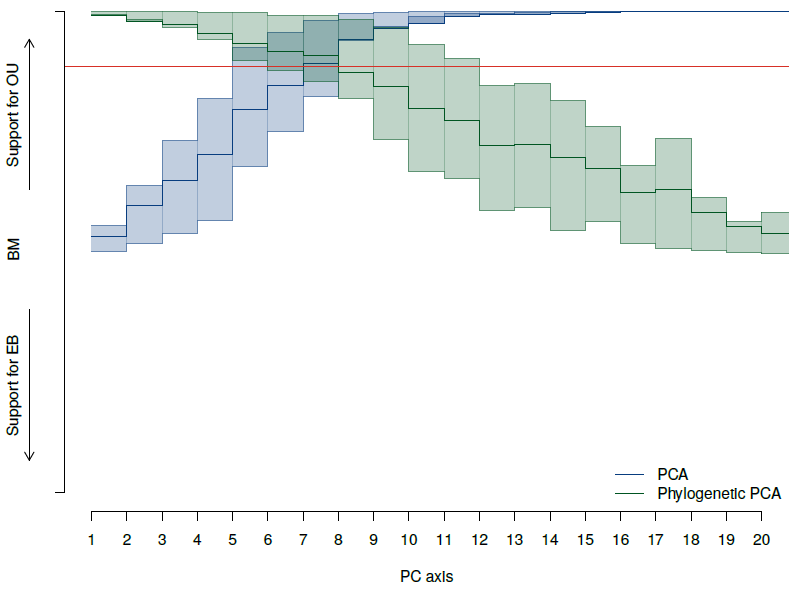
Distribution of support for BM, OU and EB models when the generating model is a uncorrelated multivariate OU model. Support for models were transformed into a linear scale by calculating an overall model support statistic: *AICw_OU_-AICW_EB_*. Thus high values support OU, low values support EB, and intermediate values near 0 indicate BM-like evolution. Models were fit to each replicated dataset for each of 20 different traits which were taken either from PC scores (blue line) or phylogenetic PC scores (green line). Shaded regions indicate the 25*^th^* and 75*^th^* quantiles of the model support statistic for 100 replicated datasets. The red line indicates the average model support statistic averaged over all 20 original trait variables. Note that standard PCA results in Akaike weights that are skewed toward EB for the first few PCs of standard PCA, while and that later PCs subsequently favor OU models. By contrast, pPCA results in Akaike weights that are skewed toward stronger support for OU models relative to the original trait variables.

**FIGURE S.2:**
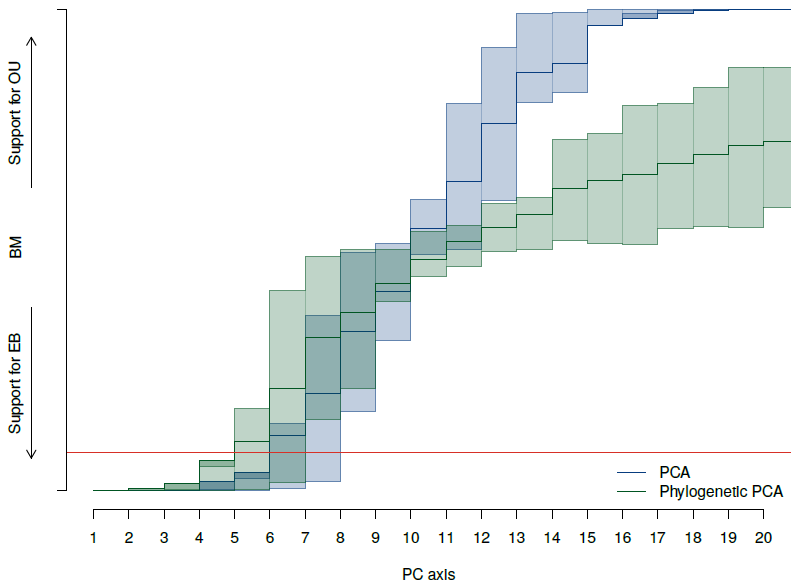
Distribution of support for BM, OU and EB models when the generating model is an uncorrelated multivariate EB model. Support for models were transformed into a linear scale by calculating an overall model support statistic: *AICw_OU_ – AICw_EB_*. Thus high values support OU, low values support EB, and intermediate values near 0 indicate BM-like evolution. Models were fit to each replicated dataset for each of 20 different traits which were taken either from PC scores (blue line) or phylogenetic PC scores (green line). Shaded regions indicate the 25*^th^* and 75*^th^* quantiles of the model support statistic for 100 replicated datasets. The red line indicates the average model support statistic averaged over all 20 original trait variables. Note that both pPCA and PCA increase support for EB models for early PC axes.

**FIGURE S.3:**
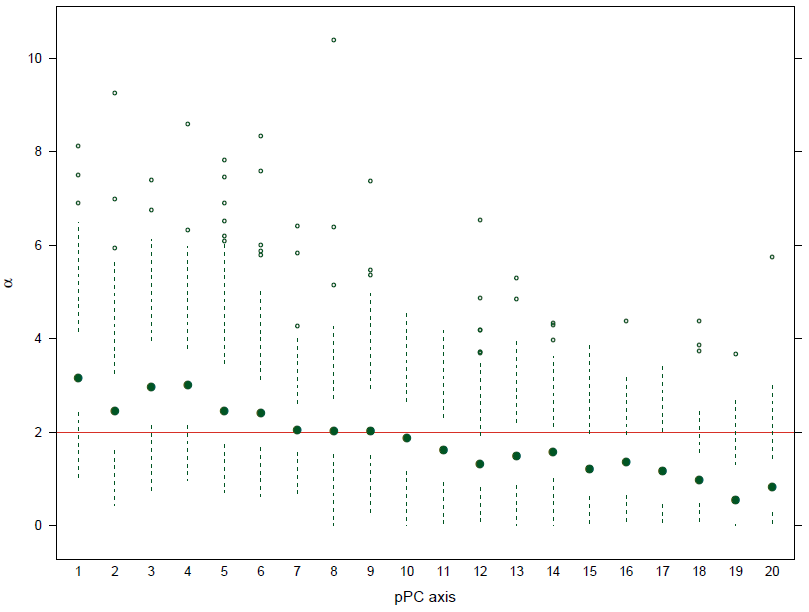
Estimated values of the *α* parameter from phylogenetic PCA when data is simulated under a correlated multivariate OU model. The simulating value *α* = 2 is depicted with the red line. The estimate of *α* is inflated in the first few pPC axes consistent with an exaggerated support for the OU model. In the last pPC axes, *α* is estimated to be very close to 0, such that the OU model is statistically indistinguishable from a BM model.

**FIGURE S.4:**
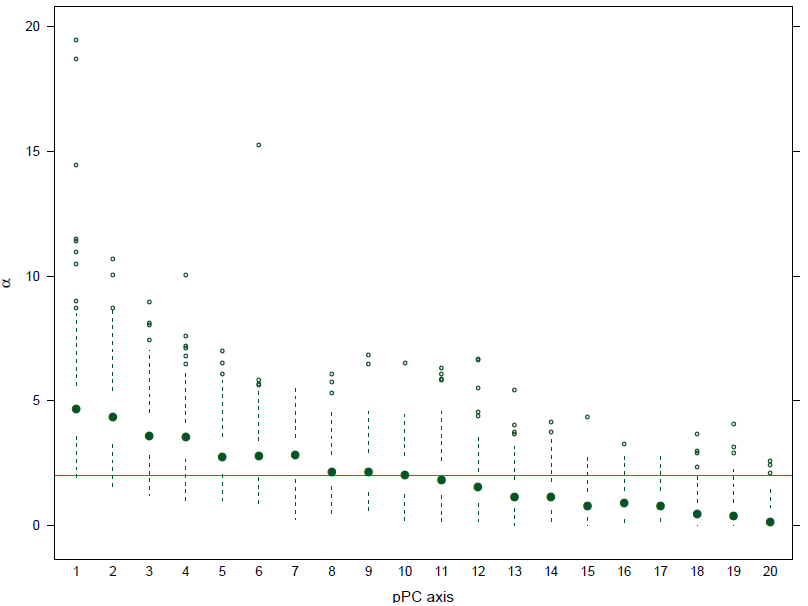
Estimated values of the *α* parameter from phylogenetic PCA when data is simulated under an uncorrelated multivariate OU model. The simulating value *α* = 2 is depicted with the red line. The estimate of *α* is inflated in the first few pPC axes consistent with an exaggerated support for the OU model. In the last pPC axes, *α* is estimated to be very close to 0, such that the OU model is statistically indistinguishable from a BM model.

**FIGURE S.5:**
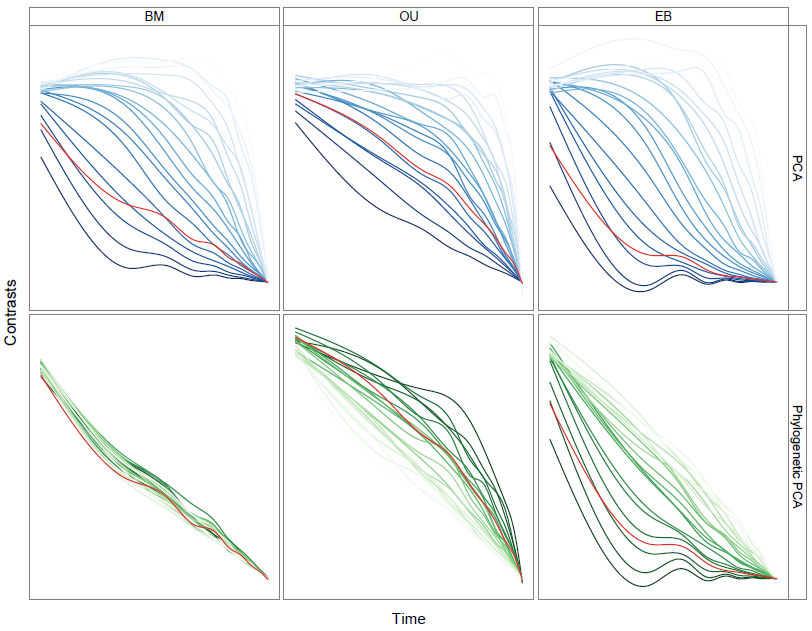
Disparity through time plots averaged across the 100 simulated datasets. The datasets were simulated under BM (left), OU (middle) or EB (right). The analyses were then performed on PC scores (top row) and pPC scores (bottom row). The average disparity through time of all 20 original trait variables is indicated by the red line. We fit a loess curve through the relative disparities for each trait/transformation/model combination. The plots are oriented so that the left side of each panel corresponds to the root of the phylogeny, with time increasing tipward to the right. The intensity of the colors are proportional to the ranking of the PC or pPC axes, stronger lines represent the first axes. As in Fig. 3, the first few axes from the PCA show a strong pattern of high disparity early in the clades’ histories with the higher components showing seemingly higher disparity towards the present. Phylogenetic PCA corrects the distortion if the generating model is multivariate BM. However, if the generating model was not BM, the first few pPC axes tend to show an exaggerated pattern of disparity relative to the original traits.

**FIGURE S.6:**
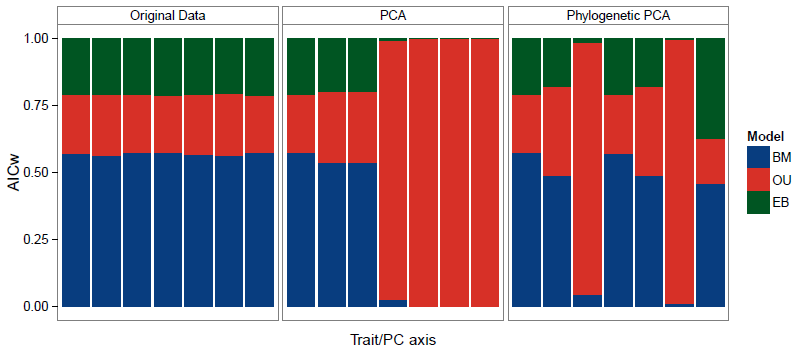
Proportion of support for BM, OU and EB models for each of the traits/PC axes from the morphological dataset of Felidae species from Slater and Van Valkenburgh (2009) and Sakamoto et al. (2010). Traits were log transformed prior to analysis. Note that all original traits and the first axes under standard and phylogenetic PCA show strong support for a BM model.

**FIGURE S.7:**
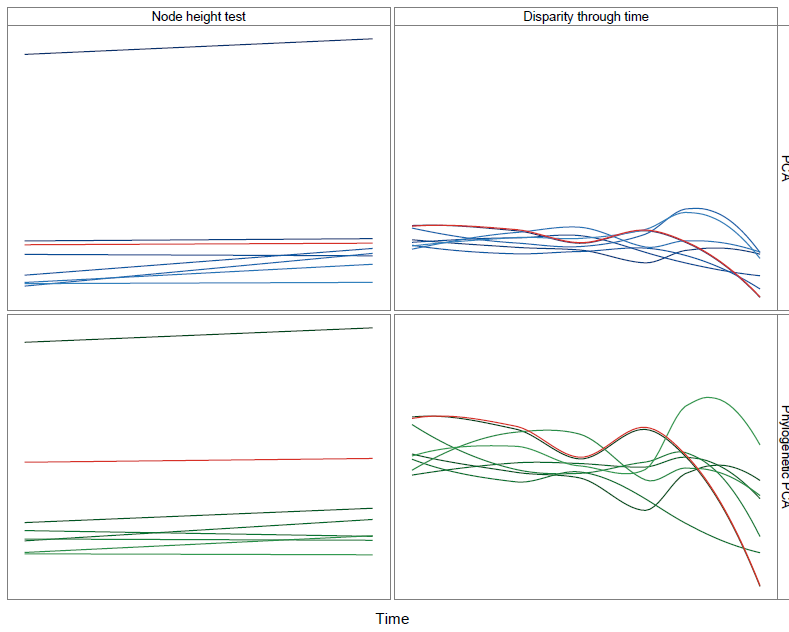
Node height test and disparity through time plots for the morphological dataset of Felidae species. Each line represents a best-fit linear model (left) or loess curve fitted (right) to the original traits, PC or pPC scores. All traits were log transformed prior to analysis. The intensity of color is proportional to the ranking of the PC or pPC axes, stronger lines represent the first axes. Left panels show the relationship between the average phylogenetic independent contrasts and the height of the node. Red lines indicate the average value for the original trait values. Right panels show disparity through time plots. The plots are oriented so that the left side of each panel corresponds to the root of the phylogeny, with time increasing tipward to the right. Compare this highly correlated dataset with only 7 traits to the larger, less correlated dataset of *Anolis* lizards (Figure S.9).

**FIGURE S.8:**
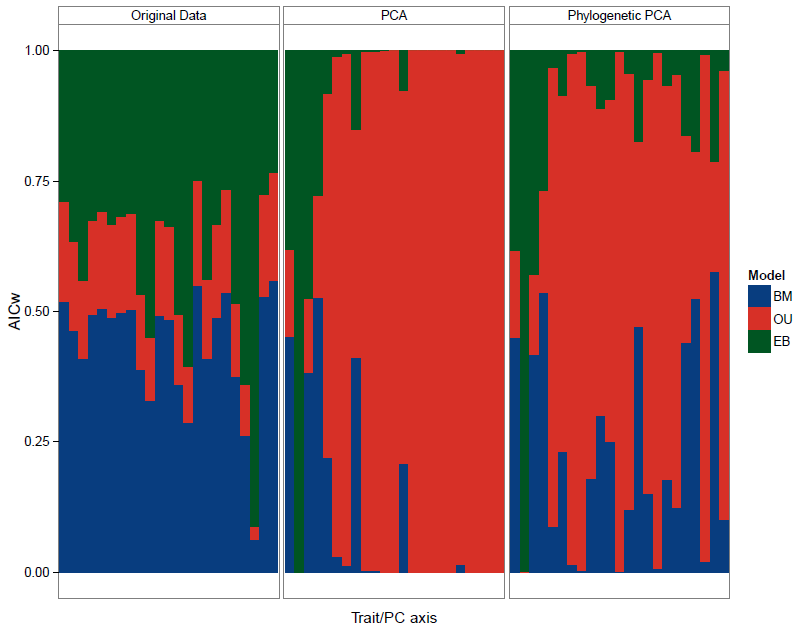
Distribution of support for BM, OU and EB models for a 23-trait morphometric dataset taken from Mahler et al. (2010). Support is measured in Akaike weights across all original trait variables (left), as well as standard PCA (middle) and pPCA (right). For both PCA and pPCA, support for the EB model appears to be concentrated in PCs 1−4, with a suggestive pattern of decreasing support across PCs 2−4.

**FIGURE S.9:**
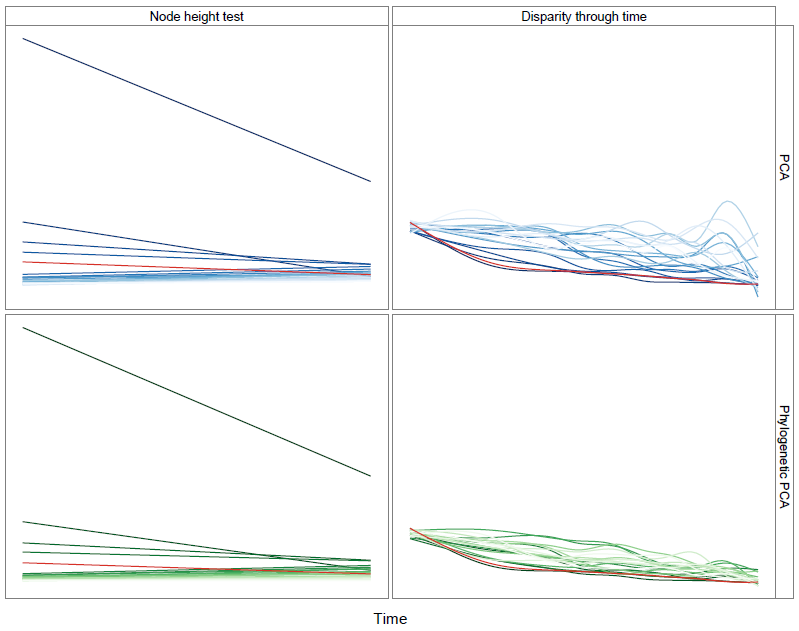
Node height test and disparity through time plots for the morphological dataset of *Anolis* lizards. Each line represents a best-fit linear model (left) or loess curve fitted (right) to the original traits, PC or pPC scores. All traits were log transformed prior to analysis. The intensity of color is proportional to the ranking of the PC or pPC axes, stronger lines represent the first axes. Left panels show the relationship between the average phylogenetic independent contrasts and the height of the node. Red lines indicate the average value for the original trait values. Right panels show disparity through time plots. The plots are oriented so that the left side of each panel corresponds to the root of the phylogeny, with time increasing tipward to the right.

